# XVCF: Exquisite Visualization of VCF Data from Genomic Experiments

**DOI:** 10.1101/2025.04.30.651450

**Authors:** Ghaida A. Almuneef, Abdulrhman Aljouie, Yahya Bokhari, Ahmed Almazroa, Mamoon Rashid

## Abstract

**Background:** High-throughput genomic analyses of germline and cancer genomes facilitate the identification of causal and actionable genetic variants. The recent advances in next-generation sequencing technology generated large-scale genomic and/or multi-omics datasets from different disease types or models. Owing to the huge volume of data coming out of genomic experiments, scientists are facing challenges in handling, manipulating, visualizing, and interpreting the data. Currently, available tools to visualize genetic variants from VCF files are not very user-friendly as most of them require knowledge of command line tools or scripts to install and run those software. Moreover, biologists or clinicians lack this knowledge of computer programming. Therefore, graphical user interface (GUI) based tools or software are needed to effectively summarize and visualize the huge volume of genomic data such as VCF data.

**Methods:** We have developed a Shiny App, interactive tool using the R programming language that utilizes other R packages like “vcfR” and “maftools” to visualize and generate quality control metrics for genetic data effectively and exquisitely. A key improvement is the addition of a user-friendly interface, providing researchers with an interactive way to explore VCF or Cancer genomics data. Our tool is powered by Shiny, making it even easier for researchers to analyze and visualize genomic variation data using a GUI. Researchers can upload their datasets and customize the analysis parameters to suit their specific research needs.

**Results:** A user-friendly interactive tool has been developed for the summarization and visualization of data related to genomic variation research. The application features an easy and friendly interface, allowing users to perform various functions such as data loading, summarization, and visualization interactively. XVCF offers an easy-to-use GUI platform to read genetic variant data (annotated or unannotated) and extract useful information such as read depth, mapping quality, genotype, quality control summary, and allele frequency from unannotated data. In the second module of XVCF, the cancer genomic data (annotated, so far supported by ANNOVAR) is analyzed using “maftools” to produce oncoplot, comparison of mutational load across different TCGA datasets, gene summary, etc. XVCF is available for free download from https://github.com/rashidma/XVCF.

**Conclusion:** XVCF Shiny web application can serve as a robust visualization and quality control GUI platform for the germline and cancer genomics dataset. We expect this tool will be immensely useful for researchers with less computational or technical knowledge. Being a shiny R package, XVCF can be installed across different operating systems and utilize different computer hardware configurations.

## 1. Introduction

Genomic analysis forms the cornerstone for unraveling the complexities of genetic variation and what they imply in various biological contexts. High-throughput sequencing technologies and platforms are resulting in voluminous data with the advent of genomic experiments like whole-genome and whole-exome sequencing and RNA-sequencing experiments [1]. Such datasets offer excellent information for genetic variation or expression patterns and become the backbone of other variant calling and measurement of expression studies [2].

Variant Call Format (VCF) represents a kind of standardized file that stores information about genetic variation detected through any sequencing experiment. The data coming from next-generation sequencing experiments could be huge in terms of several variants recorded and the file size in the computer hard disc. For example, WES data may contain 100s of thousands of genetic variants from one individual and may occupy several megabytes in the hard disc. Similarly, WGS data may contain ∼4 million genetic variants per individual and could occupy gigabytes of space. It becomes very challenging to analyze and visualize such a large amount of data coming from large cohort studies comprising 100s of individuals/patients [3]. Thus, efficient algorithms and tools are very critical and demand time for visualization and summarization of large genomic datasets comprised of VCF files.

Among the existing VCF filtering and visualization tools are: VCFtools [3], GEMINI [4], BrowseVCF [5], VCF.Filter [6], VCF-Miner [7], VCF-Server [8], VIVA [9], vcfR [10], vcfView [11], and IGV [12]. We only included open-source technologies published in peer-reviewed papers. While these provide powerful functionalities, they also pose significant challenges for the end-users to install and run these software provided the little computer knowledge and resources available. Most of the above-mentioned tools are filtering tools, so we will not describe them in detail since our focus here is on the visualization aspect.

Starting with an excellent illustration tool named VIVA [9], which is a command-line utility and Jupyter notebook-based tool that evaluates and shares genomic data from VCF files for variant analysis and quality control of sequencing projects. VIVA is distinguished by its simplified and visually appealing working environment for reviewing and sharing genomic data and its visualization approaches for producing publication-quality graphic visuals with variant analysis. This allows for exploring phenotype-genotype associations, batch-effect assessment on coverage, and differential variant incidences across samples. However, the command-line interface must be very daunting for researchers who are not at all computational, whereas the use of Jupyter Notebooks restricts access to VIVA among more users.

Second, we have vcfR [10]: An R package useful in browsing and parsing VCF files, for detailed analysis. In addition, vcfR is built over the powerful statistic and visualization libraries of R and uses the functions implemented there, so it could be even faster at processing data, particularly given a direct integration of sequence (in FASTA) and annotation (in GFF) data for detailed genomic region visualization. The package also enables exporting data into other R genetics packages and implements all computationally intense functions in C++ to improve performance. However, the steep learning curve associated with R programming and the complexity in the layout of VCF files may render the tool nonuser-friendly or less accessible for some users.

On the other hand, two novel tools were developed specifically to facilitate the visualization of genomic data: VcfView [11] and IGV [12]. VcfView is an interactive tool developed for comparing somatic mutation calls made from cancer sequencing data. The researchers developed a graphical user interface that allows visualizations of VCF Files the dashboard allows interactive filtering and tracking of the impacts of analytical decisions on the mutant allele frequency spectrum, mutational signatures, and the aggregated somatic variants of a gene. Nevertheless, this tool cannot be universally applied. VcfView is limited to matched cancer/normal VCF files and the use of VCFs generated only by Mutect2, which reduces the tool’s universality.

The Integrative Genomics Viewer (IGV) is a graphical viewer that displays multiple alignments, variants, and various genomic elements. It was intended to be easy enough to use that exploring large-scale datasets is as easy and intuitive as using Google Maps. IGV is a perfect tool to integrate multiple sequence read alignments with a panel of other types of data and ease access to these data on the cloud, making it a powerful tool for a shared data repository. IGV, on the other hand, is also constrained by the abstractions it uses – with a greater abstraction level, it is difficult for users to interact with the specific elements of the app. Despite being competent, IGV may lack some of the more expert tools and functionality for more extensive and more critical projects.

Though each of these tools has advantages over the others, there are also certain limitations attached that limit their widespread use. The complexity of genomic data and the diverse requirements of research projects bring out the need to go in-depth into the sensitivity of all the strengths and weaknesses of each tool. Here, we will discuss the development of our offered solution: A GUI-based Shiny app that can help better visualize genomic data and generate intuitive quality control plots to make the research process easier and more effective.

The idea is to modify the way visualization and summarization of genomic data sets stored in VCF files and annotated VCF files in tabular format are done. Our app democratizes the space of genetic data analysis by offering users an intuitive and easy-to-understand platform on which to explore and make sense of their data more efficiently. Our tool takes a step further in neutralizing the existing barriers related to VCF file analysis by fostering user-friendliness and easy access. It further helps to foster a more inclusive and smoother research process.

## 2. Methodology

### 2.1 Development of Shiny Web Application to Visualize Genetic Variant Data

To visualize large-scale high-throughput genetic variation data, coming out of genome and/or transcriptome sequencing projects, there has always been a need for an easy-to-use, lightweight, and GUI-based tool. We addressed this gap and made an effort to develop such a GUI-based tool called “XVCF” using the Shiny application in the R environment (R version 4.2.1). We integrated the two most widely used R packages named “vcfR” [10] and “maftools” [13] for reading, generating quality control plots, and visualizing the most available variant call format (VCF) file(s) containing genomic variant data.

### 2.2 Design and Features

Through the use of the Shiny server, we designed an interface that offers easy-to-use functionalities such as uploading data, summarizing VCF files, and producing high-quality plots of important parameters The interface is dual accepting both i) VCF file and ii) annotated VCF file in tabular format (ANNOVAR is currently supported) separately. The first module accepts VCF files and processes through our proprietary code as well as through various functions available in vcfR package. This interface includes tools for visualization where users can upload their datasets and check summaries of their VCF files. Additionally, users can examine parameters from VCF files like read depth, genotype quality, allele frequencies, genotype plots, and the distribution of quality metrics in different forms (bar plot, boxplot, heatmap, Histogram, and Violin plot). The second module of XVCF accepts ANNOVAR output in tabular format which is converted to MAF format using “maftools” [13]. We ask the user to enter the number of samples first before uploading the ANNOVAR file. There is an option for the user that allows them to apply some filtrations on allele frequency (AF) to enrich somatic mutation. The tool showcases key plots like the ANNOVAR Summary Plot and oncoplot for the top ten mutated genes, alongside titv summary plots, lollipop plot, the rainfall plot, Compare mutational load against TCGA cohorts, and Oncogenic Pathways plot. Also, generates summaries such as sample summary, gene summary, and clinical summary.

The overall design and features of the XVCF tool are depicted as a schematic in Figure 1.

**Figure 1.**
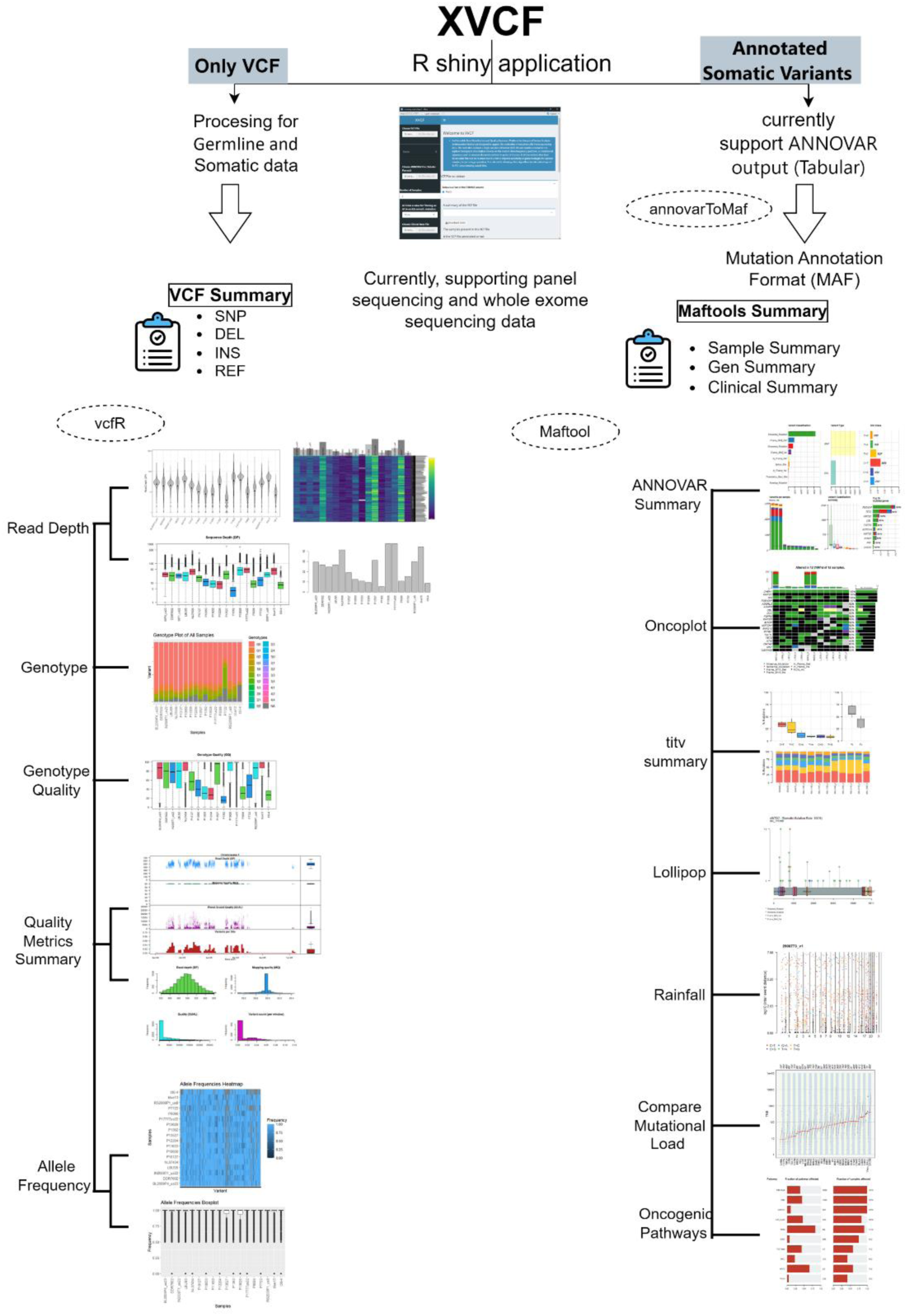
Schematic diagram showing the overall design and features of the XVCF app.

### 2.3 VCF File Structure and Pre-processing

The Variant Call Format (VCF) serves as a file format, for storing variations like SNPs, insertions, deletions, and structural changes. A VCF file consists of meta-information lines, a header line, and data lines containing information about a position in the genome.

The Meta information lines that begin with ## and define the abbreviations and metadata used in the file. The data section is presented in a format where each row represents a variant along with details about it in columns.

The initial eight mandatory columns in the data section provide information on the chromosome (CHROM) starting position of the variant (POS) an identifier (ID) the reference allele (REF) alternative allele(s) (ALT) Phred scaled quality score (QUAL) a flag indicating successful or failed control checks (FILTER) and annotations specific, to the variant (INFO). If multiple samples are included additional columns contain genotype details (FORMAT) and corresponding sample information. This intricate arrangement allows for data inclusion. It requires careful handling to manage the variability associated with each variant.

Our user-friendly Shiny web application simplifies the process of loading and visualizing VCF files. Users have the option to upload uncompressed or compressed (*.gz) VCF files, through a file selection interface, then the system preprocesses the VCF file to extract data for visualization purposes. The same applies to the Cancer genomics data and the clinical data. Users can choose from a range of plotting options using a sidebar to generate plots ensuring they are available for immediate download. This feature enables swift exploration of data.

### 2.4 Implementation Details

Creating an R/Shiny application entails setting up a coding environment with the package, in R, which allows for the creation of web applications featuring engaging user interfaces.

#### 2.4.1 Environment Setup

Creating an R/ app requires having an R environment, which you can download from CRAN and install on operating systems, like Windows, macOS, and UNIX. While it’s not a must it’s recommended to use RStudio, an IDE designed for R programming when developing apps. Also, make sure to install the package from CRAN.

#### 2.4.2 Application Structure

A typical R/Shiny application consists of two R scripts, “ui.R” and “server.R”, which are usually placed in the directory of the application. The ui.R script is responsible, for defining the elements of the application by setting up its layout and appearance using “Shiny/shinydashboard” layout functions like: “dashboardPage”, “dashboardSidebar”, and “sidebarMenu”. Within ui.R, various input widgets can be created to gather user inputs such as uploading files pasting text, or adjusting parameters. These user inputs captured by “ui.R” are then sent to the server side for processing by “server.R”. The results generated by “server.R” are subsequently presented as plots, tables, etc. at designated spots determined in “ui.R” in the user interface. To simplify things for applications in R/Shiny you can combine the code from both “ui.R” and “server.R”, into an R script.

#### 2.4.3 Running the Application

The “shinyApp()/Run App” function, in the package allows for the initiation of an R/ app by providing the directory path of the application. You can Run the R/ app in RStudios viewer panel, a new window, or your system’s default web browser. It’s suggested to run the R/ app in a web browser. Any changes made to the appearance or features of the R/ app, through code updates, can be seen immediately without refreshing the web browser.

#### 2.4.4 Designing the UI

When creating an R/ application you start by designing the user interface using ui.R. The dashboard application’s user interface is constructed using the “shinydashboard” framework, where the root element is “dashboardPage”. The header component which is created using the “dashboardHeader()” contains the title “XVCF”. The sidebar navigation is developed by the “dashboardSidebar” where the menu is organized in a hierarchical manner starting with the file input component for the VCF file. The main menu is developed by the “sidebarMenu” with the root menu item named “Home” (tabName = “home”) which contains several submenus with names corresponding to the types of visualizations that can be chosen. Other features are included in the form of “shinyjs()” for improving the interactivity of the application and numeric input to specify the sample. The sidebar also contains a select input for filtering by AF threshold and file input for clinical data. The body section is developed with the help of “dashboardBody”, and it also includes “tabItems()” to organize the sections of the dashboard. The dashboard can display images, tables, and text. By default, the sidebar is positioned on the side of the dashboard but can be switched to the right side by adjusting the position parameter.

##### 2.4.5 Reactive Programming Model

Shiny apps depend on a programming approach enabling instant user interface changes in response to user actions. Reactive expressions “reactiveVal” and “observeEvent” functions play a role in handling reactivity. For example, when a user uploads a VCF file the server function processes the file. Updates the UI with summaries and visual displays smoothly removing the necessity for page reloads.

#### 2.4.6 Integration of vcfR and maftools

The vcfR [10] package is used to manage and manipulate VCF files with features, like “read.vcfR” for reading VCF files, “extract.gt” for retrieving genotype information, and “heatmap_bp” for displaying read depth. The” maftools” [13] package streamlines the analysis and display of mutation annotation data offering functions such as “annovarToMaf” to convert ANNOVAR results to MAF format “read.maf”, for reading MAF files and visualization tools like “plotmafSummary”, “oncoplot”, “titv”, “lollipopPlot” and “rainfallPlot” for creating mutation related visualizations.

#### 2.4.7 Application Deployment

The Shiny application is configured on a server system to provide access to researchers. Setting up involves adjusting the server environment and hosting the application enabling users to reach it through a web interface.

### 2.5 Technical Aspects

- **Platform Compatibility:** The software is created exclusively in R guaranteeing performance on operating systems. It has been trialed on RStudio with base R 4.2.1 and Shiny 1.8.0.
- **Performance enhancement**: We’ve enhanced the performance of XVCF using existing optimized R packages such as “vcfR” version 1.15.0 and “maftools” version 2.12.0.
- **User-Friendly Design:** The framework helps users effortlessly access the application, on devices and operating systems without the hassle of intricate setup. You can get the app, from GitHub at “https;//github.com/”. All data processing happens in R to ensure it works on any platform. There’s also a guide based on a dataset available on GitHub.

## 3. Results and Discussion

### 3.1 The Shiny Application Interface Overview

In this study, we have developed a user web application that aims to visualize genomic variant data and ensure its quality. The interface of the application is titled “An Extensible Data Visualization and Quality Assurance Platform, for Integrated Variant Analysis … etc.” emphasizing its nature (Figure. 2). Researchers can utilize this tool to evaluate mutation calls from sequencing data by inputting a Variant Call Format (VCF) file. Through this platform, users can explore the impact of choices on parameters such as mutant allele frequency spectrum, mutational signatures, and annotated somatic variants in specific genes. Moreover, researchers have the opportunity to review variants that did not pass caller filters, which helps improve sensitivity and assists in refining sample or sequencing preparation strategies.

**Figure 2.**
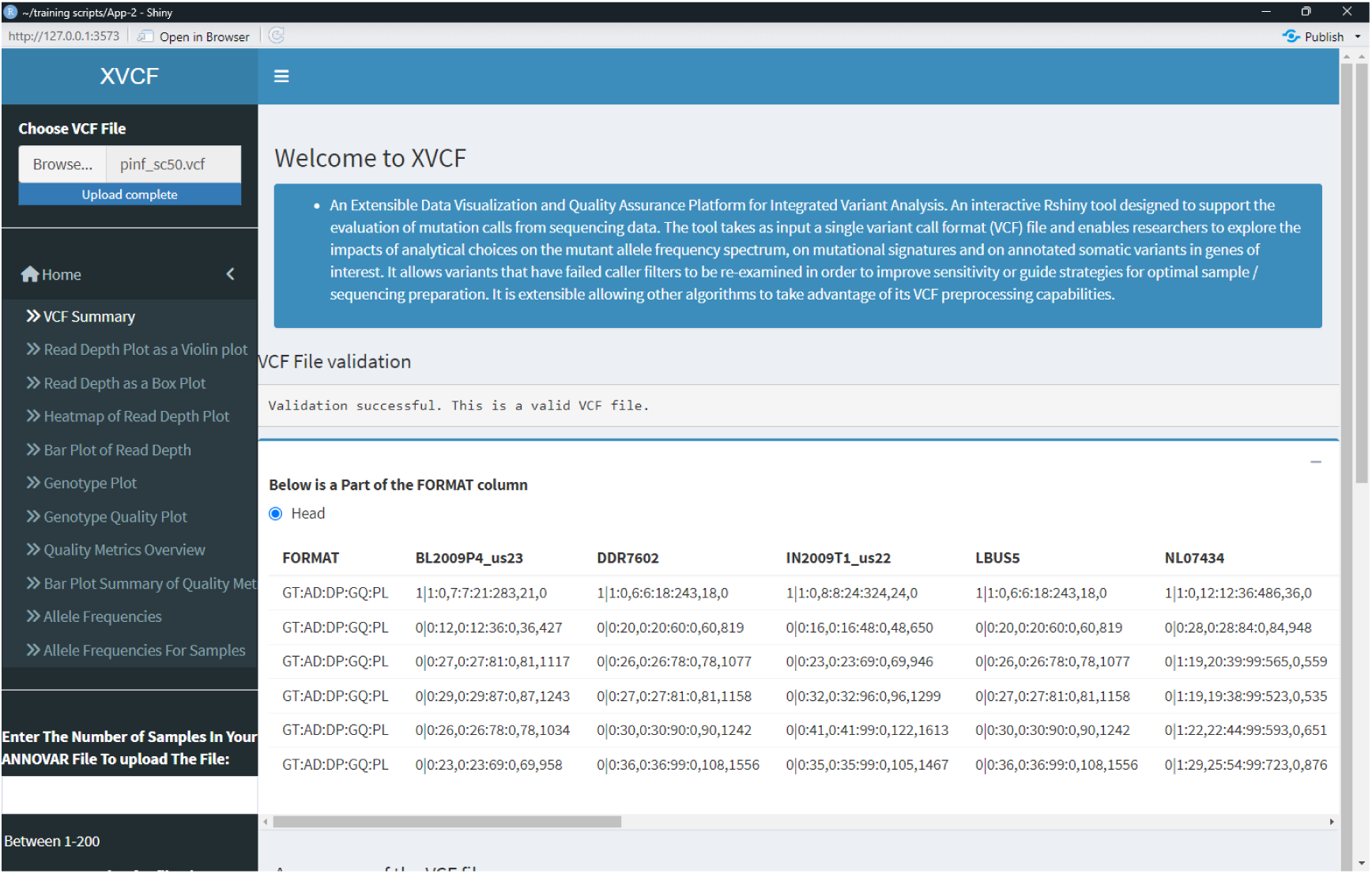
The shiny app interface.

### 3.2 Summary of the VCF File

When the VCF file is uploaded to the application the users receive a detailed summary that includes several important aspects. Starting with a Validation of the VCF file format states that the VCF file meets certain standards to ensure compliance with specifications. Followed by a small display containing the first 6 lines (Header)of the VCF FORMAT column. After that, there is a table containing information such as the Reference details to give information about the reference genome used for the file. As well as details about contigs to provide information regarding the contigs of the Chromosome listed in the VCF file. Also, it contains the number of samples and sample names to highlight the sample count and their names as shown in the file. Then a Variants Overview to cover the types of variations and their count including single nucleotide polymorphisms (SNPs), insertions, deletions, and other genetic variants. Lastly, the phasing and annotations status provides information on whether the VCF file is phased or un-phased and if it’s annotated or not.

This summary feature is very important as it helps determine whether the VCF file is well structured and can be analyzed, or not hence ensuring data quality.

### 3.3 Interactive Plotting Functions

The software allows users to easily upload datasets, in VCF and annotated VCF files in tabular formats along with clinical data files in Excel format making data input and visualization simpler. Users have the option to customize some settings according to their research requirements and queries with the freedom to adjust visualization preferences for a personalized visualization experience.

Users have the option to choose plots, from the sidebar on the left side to see displays generated through the “vcfR” and “maftools” packages starting with:

#### 3.3.1 Visualization of Read Depth

The read depth is the frequency of a base appearing in all the sequences read during sequencing. Scientists can be certain in identifying a base through ‘base calling’ with increased read depth. We can provide the users with different plots for the read depth (DP) extracted from the VCF FORMAT column highlighting the median, and any potential outliers in the data. This visualization is helpful for identifying samples with low coverage. The read-depth plots are created by different code blocks utilizing the data extracted from the VCF file with the function “vcfR:: extract.gt()”. Four different types of visualization have been created with read depth data such as violin plot (Figure 3A), box plot (Figure 3B), heatmap (Figure 3C), and bar plot (Figure 3D).

**Figure 3.**
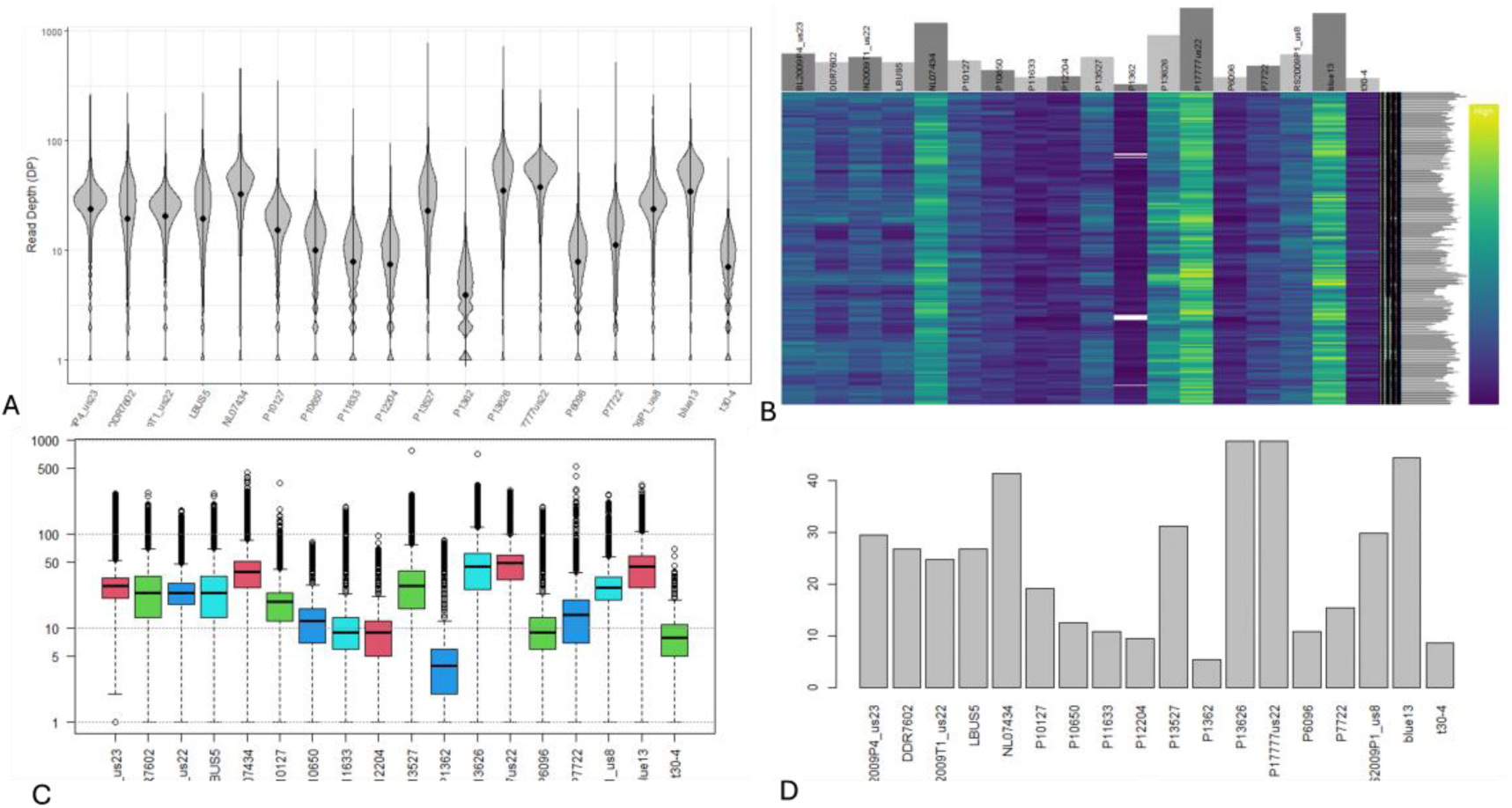
The Visualization of the Read Depth in Different Ways (A) Read Depth as a Violin Plot for18 samples, (B) Read Depth as a Box Plot for18 samples (C) Heatmap of Read Depth Plot, Each column is a sample, and each row is a variant. Each cell’s color represents each variant’s read depth (DP). (D) Bar Plot of the actual values of Read Depth where the x-axis presents the samples and the Y-axis shows the DP. (All the images were created from the pinfsc50 data set).

#### 3.3.2 Genotype Information and Quality Visualizations

The term genotype describes the collection of alleles that an organism possesses for a trait. Alleles are gene variations and genotypes in a VCF file are commonly denoted, like 0|0 (homozygous for reference), 0|1 (heterozygous for alternate), or 1|1 (homozygous for alternate). We provide the user with two options regarding genotype information as mentioned below:

1. Genotype Visualization: In terms of genotype visualization, this graphical representation exhibits the types of genotypes in the VCF file. The horizontal axis represents samples while the vertical axis represents variants assisting in grasping how homozygous and heterozygous variants are distributed within the dataset (Figure 4A). Both Genotype and Genotype Quality plot was obtained from VCF genotype data using the “extract.gt” function, but with different arguments, we select: element = “GT” to extract the genotype from the extracted matrix.
2. Plot of Genotype Quality: We have a plot visualizing genotype quality (GQ) scores to assess the reliability of genotype calls (Figure 4B). High GQ scores indicate predictions and this graph assists, in pinpointing variations with possibly questionable genotype quality.

**Figure 4.**
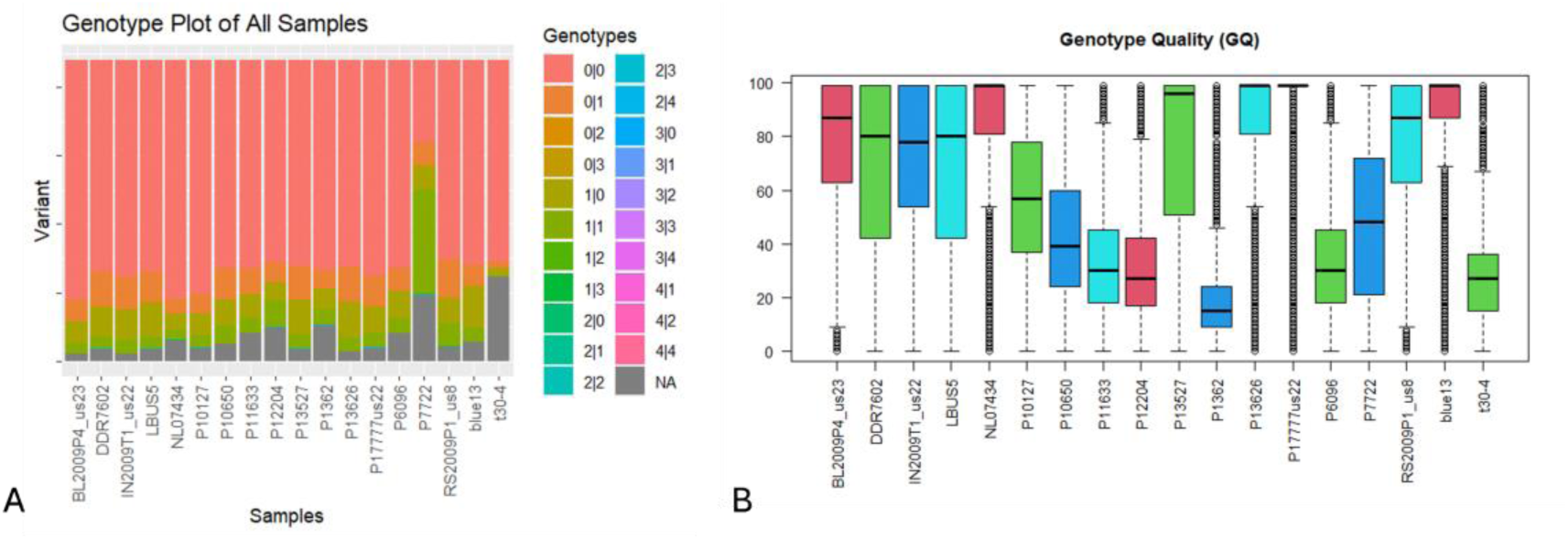
We present two use cases for Genotype. In the first use case, A. shows the genotype plot of all samples in the VCF with their unique variant, where the x-axis presents the samples and the Y-axis shows the GT (usually the VCF stores the genotype as: “0|0, 0|1, 1\1” however the data set we used “pinfsc50” contain more options) B. shows the genotype Quality plot of all samples in the VCF, where the x-axis presents the samples and the Y-axis shows the mean GQ value for each sample. (All the images were created from the pinfsc50 data set).

#### 3.3.3 Summaries on Margins and Quality Metrics

Variant calling programs usually need to undergo post hoc quality filtering. This becomes crucial in standard systems, with limited curated data availability [14]. The VCFR function “chromoqc()” was used to generate (Figure.5) from the VCF file only. However, it usually requires three files: a FASTA sequence file, a GFF annotation file, and a VCF file for extra information about annotations, and analysis of GC content. The plots in this section were as:

i. Brief Summaries on Margins (Multi-Layered Summary of Quality Metrics): These visual summaries present an overview of the data extracted from the VCF file including counts per section and scatter plots showing quality measures like variant incidence, mapping quality, read depth, and phred-scaled quality (QUAL). These summaries give a top-level view of data quality and its distribution (Figure. 5A).
ii. Graph Illustrating Quality Metrics Distribution(Histogram Summary of Quality Metrics): This visualization showcases the spread of quality metrics obtained from the VCF data, such as variant incidence, mapping quality, read depth, and phred-scaled quality (QUAL), as bar plots (Figure. 5B). It aids researchers in evaluating the sequencing data quality. Summarized through “proc.chromR()” function.

**Figure 5.**
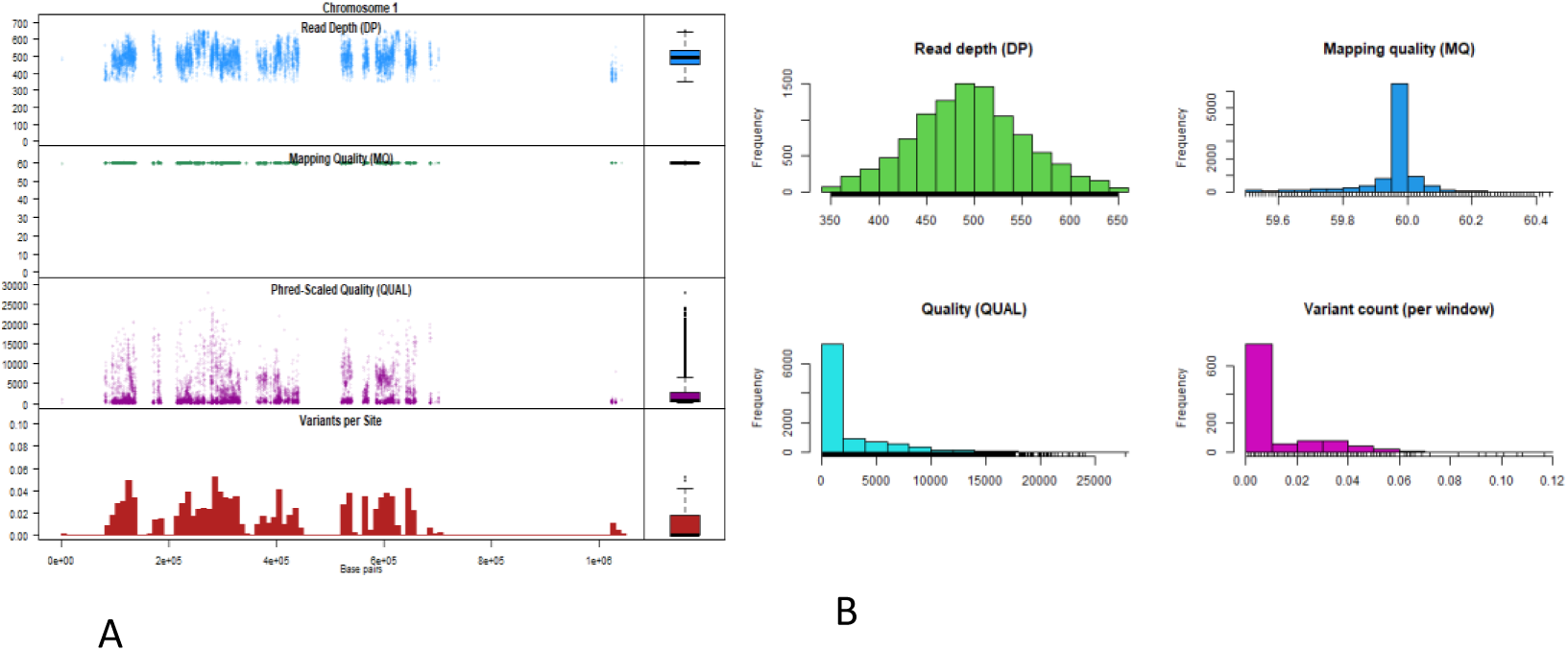
Chromoqc plot showing raw variant call format (VCF) data for one super-contig (only one chromosome) in the pinfsc50 data set. (A.) The lowest panel represents one of the variant incidences. Above this are three-dot plots of phred-scaled quality (QUAL), Continuing up the plot are mapping quality (MQ) and read depth (DP). (B.) It shows the same quality metrics variant incidence, mapping quality, read depth, and phred-scaled quality (QUAL), but as bar plots.

#### 3.3.4 Plots on Allele Frequency

Allele Frequency (AF) usually indicates how often a specific gene variant occurs in a group of individuals. It shows the percentage of that allele in the population, ranging from 0 to 1. When there are alleles at a genetic location, the total of all allele frequencies for those alleles at that location should equal to 1.

The plots utilize the “AD_frequency” function, from the “vcfR” package to generate allele frequencies based on matrices of allelic depths (AD). The user has two options for this plot either as:

i. Heatmap Displaying Allele Frequencies: This heatmap exhibits allele frequencies across samples and variants. It serves as a tool for representing allele frequency distribution and identifying potential rare or common variants (Figure. 6A).
ii. Boxplot Showing Allele Frequencies: Similar, to the heatmap this box plot displays allele frequencies offering a depiction of how allele frequencies are distributed in the dataset, however, it will represent the mean value of AF in the samples (Figure 6B).

**Figure 6.**
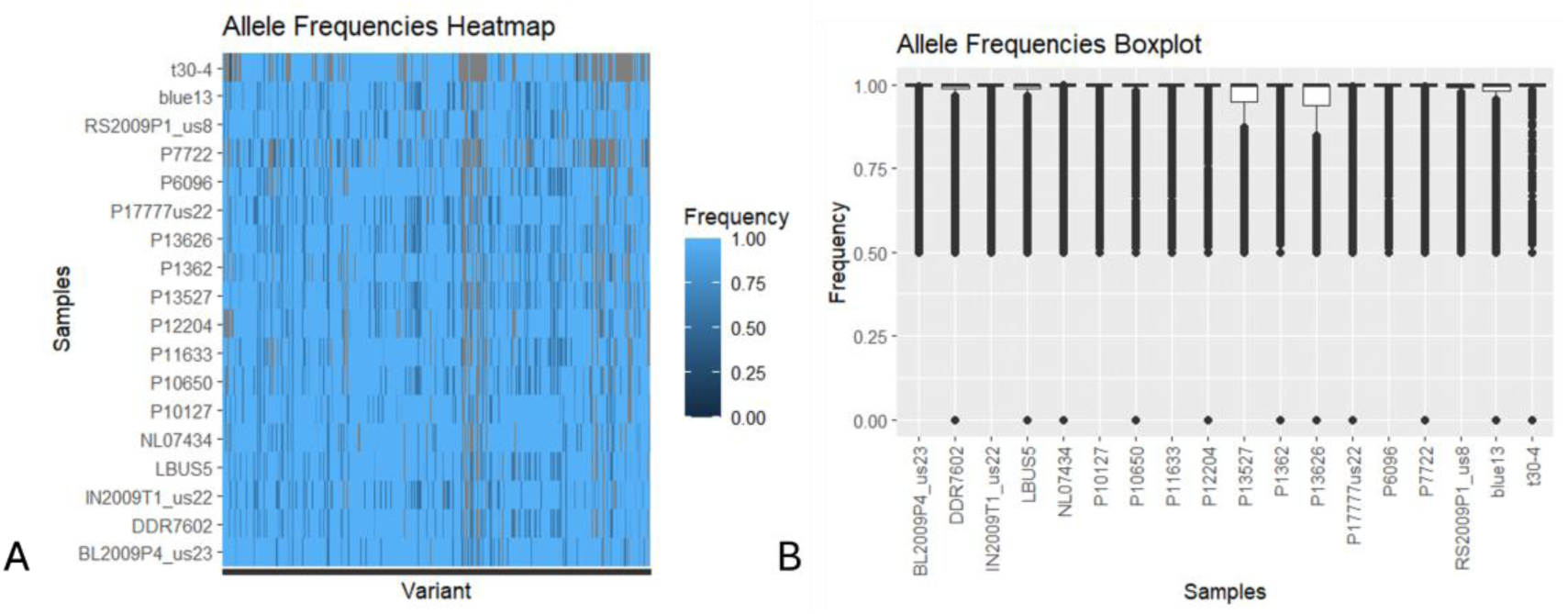
We present two use cases of allele frequency plots. In the first use case (a) we visualize variants in a heatmap of allele frequency for a set of 18 samples. Allele frequency positions are stored in rows each color represents a different value and individual samples are stored in columns. In the second use case (b) we visualize the allele frequency as a box plot where the X axis shows the samples which are 18 samples and the Y axis represents the average value of the allele frequency for each sample.

### 3.4 Cancer Genomics Data Visualizations Using maftools

The second module of XVCF accepts ANNOVAR output in tabular format for annotating variants which is converted to MAF format using “maftools::annovarToMaf”. In this section, the user can get several different summaries and plots with more flexibility to choose some preferences like filtering options or choosing the gene of interest to be plotted or by samples.

#### 3.4.1 Summary of Genes, Samples, and Metadata

This feature gives researchers a snapshot of gene, sample, and metadata details using the function “maftool:: getSampleSummary” for sample summary which contains the type of mutation (Missense, Nonsense, Frameshift Del, Frameshift Ins, In Frame Del, In Frame Ins …) and the total number of mutations in each sample, “getGeneSummary” to Show gene summary that similarly includes the type of mutation and the number of muted sample along with the total number of mutations in each gen, and “getClinicalData” for displaying clinical data associated with samples (optional). All of this information is derived from Cancer genomics data offering a summary of the dataset.

#### 3.4.2 Comprehensive Summary Plot from ANNOVAR

This detailed plot created with the function “plotmafsummary” visually represents mutation annotation data providing insights into the mutation landscape within the dataset as shown in Figure (7-A) to reveal variant classification per mutations, samples, type, SNV class, top 10 muted genes.

**Figure 7.**
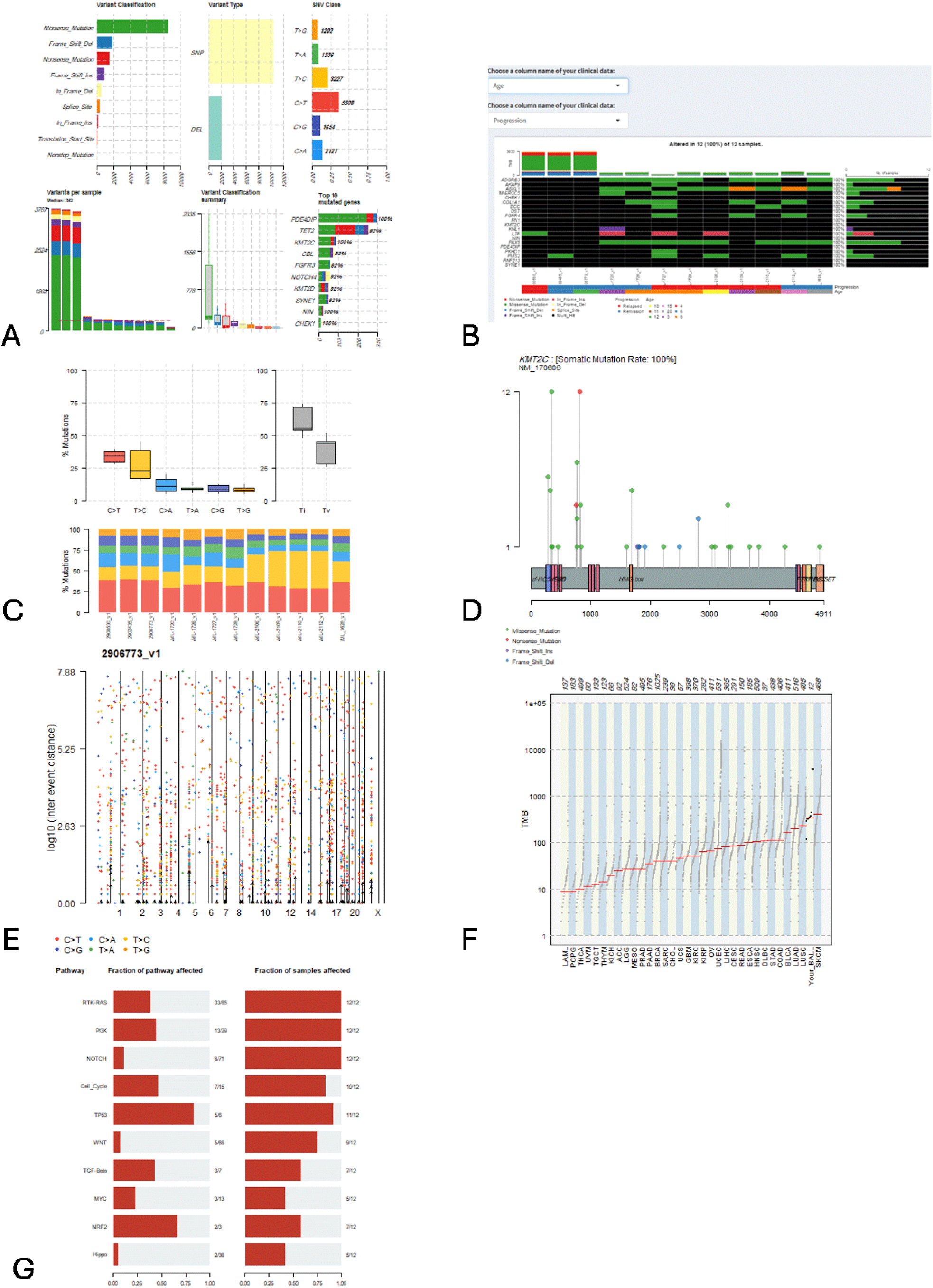
A) ANNOVAR Summary Plot. B) oncoplot for top ten mutated genes. C) plot titv summary. D) lollipop plot. E) The rainfall plot of the sample 2906773_v1 shows the distribution of the mutations across the genome. F) Compare mutational load against TCGA cohorts. G) Oncogenic Pathways plot.

#### 3.4.3 Heatmap Showing Top Ten Mutated Genes

The “oncoplot” function generates a heatmap displaying the ten frequently mutated genes highlighting significant genetic changes in the dataset. It has an option if the user uploads a clinical data file to display this information in the plot (such as Age, Gender, stage…etc.) as demonstrated in Figure (7-B).

#### 3.4.4 Transition/Transversion (Ti/Tv) Ratio Plot

This plot illustrates the Ti/Tv ratio, a metric that indicates the balance between transition mutations (purine to purine or pyrimidine to pyrimidine) and transversion mutations (purine to pyrimidine or vice versa). This plot was done using the function “plotTiTv”. It is presented in Figure (7-C).

#### 3.4.5 Visualization of Mutations with Lollipop Plot

The lollipop plot offers an in-depth view of mutations, in specific genes of interest using the “lollipopPlot” function. By showing their position and frequency along the gene structure in Figure (7-D). Users can choose the gene of entreat from a menu item extracted from the file.

#### 3.4.6 Rainfall Distribution Visualization

The rainfall chart shows how mutations are spread across the genome, helping to pinpoint areas with mutation rates or clusters of mutations. In Figure (7-E) we can see the mutation as a rain drop among the chromosome number for each sample separately and the user can choose the sample name. The plot was generated by “rainfallPlot” function.

#### 3.4.7 Comparison of Mutational Load with TCGA Cohorts

This chart is generated using the “tcgaCompare” function to compare the amount of mutations in the dataset with that of The Cancer Genome Atlas (TCGA) groups providing a reference point for the mutation levels seen in the study on Figure (7-F). There is an option for the user to enter their own data set name to be printed in the plot next to the other studies.

#### 3.4.8 Highlighting Impact on Oncogenic Pathways by Mutations

This visual representation indicates which cancer-causing pathways are influenced by the observed mutations giving insights into the effects of these genetic changes as represented in Figure (7-G). We used the “OncogenicPathways” function to create this plot.

## 4. Conclusion

In conclusion, the Shiny web application created in this research offers a user-friendly platform for analyzing and visualizing genomic variation data. By utilizing established R packages like “vcfR” and “maftools” this tool enables researchers to effectively handle VCF files and Cancer genomics data making it a valuable asset, for research. This tool is capable of creating interactive plots/summaries and thus enables the user to fully investigate the details of genomic factors in the form of plots and tables, which allows this data to be displayed in a visual way that is easy to understand and comprehend. Thus greatly improving the efficiency and reliability of the data analysis. Furthermore, the way the design is set up and how it responds to user actions ensures a quick user experience making it possible for updates to happen in time and for users to interact with the data easily.

Nevertheless, there are areas where enhancements can be made. Improving how efficiently the app processes information especially when dealing with amounts of data well as broadening the scope of supported file formats could increase its usefulness. Moreover integrating ways to visualize data and adding interactive elements could offer deeper insights, into intricate genetic interactions.

In general, this Shiny web application marks an advancement in simplifying and streamlining genomic data analysis with room, for growth and enhancement to cater to the changing demands of the scientific community.

## Notes

### Competing Interest Statement

The authors have declared no competing interest.

### Funding

- King Abdullah International Medical Research Center, Riyadh, Saudi Arabia: NRR24/086/10

